# Live-Cell Tracking of γ-H2AX Kinetics Reveals the Distinct Modes of ATM and DNA-PK in Immediate Response to DNA Damage

**DOI:** 10.1101/2022.10.09.511457

**Authors:** Watanya Trakarnphornsombat, Hiroshi Kimura

## Abstract

DNA double-strand break (DSB) is a serious form of DNA damage that can cause genetic mutation. On the induction of DSBs, histone H2AX becomes phosphorylated by kinases, including ataxia telangiectasia-mutated (ATM), ataxia telangiectasia and Rad3-related (ATR), and DNA-dependent protein kinase (DNA-PK). Phosphorylated H2AX (γ-H2AX) can be a platform to recruit DNA repair machinery. Here we analyzed the immediate early kinetics of γ-H2AX upon laser-induced DNA damage in ATM-proficient and -deficient living cells by using fluorescently labeled antigen-binding fragments specific for γ-H2AX. The accumulation kinetics of γ-H2AX were similar in both ATM-proficient and -deficient cells. However, γ-H2AX accumulation was delayed when the cells were treated with a DNA-PK inhibitor, suggesting that DNA-PK rapidly phosphorylates H2AX at DSB sites. Ku80, a DNA-PK subunit, diffused freely in the nucleus without DNA damage, whereas ATM repeatedly bound to and dissociated from chromatin. The H2AX phosphorylation activity of ATM, but not DNA-PK, depended on a histone H4K16 acetyltransferase, males absent on the first (MOF). These results suggest distinct actions of ATM and DNA-PK that plays a primary role in immediate early γ-H2AX accumulation.

## INTRODUCTION

DNA double-strand break (DSB) is one of the most deleterious forms of DNA damage because even a single DSB can activate the DNA damage checkpoint that delays cell-cycle progression (van den Berg et al., 2018) and triggers cell death (Rich et al., 2000). These DSBs are naturally occurring with only 10–50 events per cell per day (Gospodinov and Ugrinova, 2019; Tubbs and Nussenzweig, 2017; Vilenchik and Knudson, 2003), but they threaten genomic integrity that is essential for regulation of cellular homeostasis and the maintenance of genetic information. If the DSB repair process is not properly performed, various types of mutations can arise, which may eventually lead to diseases, such as cancer (Negrini et al., 2010; Tubbs and Nussenzweig, 2017) and aging (Tian et al., 2019).

In the DSB repair response, protein kinases belonging to the phosphatidylinositol 3-kinase-related kinase (PIKK) family, including ataxia telangiectasia-mutated (ATM), ataxia telangiectasia and Rad3-related (ATR), and DNA-dependent protein kinase (DNA-PK), have critical roles. These kinases phosphorylate various proteins involved in DSB repair and the histone H2A variant H2AX at serine 139 (Blackford and Jackson, 2017). Serine 139-phosphorylated H2AX, called γ-H2AX (Rogakou et al., 1998), facilitates the concentration of DNA damage repair machinery (Celeste et al., 2003) and serves as a DNA damage signal (Hunt et al., 2013). Large-scale proteomic analysis has identified >700 proteins that are phosphorylated by ATM and ATR upon ionizing radiation, demonstrating that multiple protein networks are involved in DNA damage repair and signaling processes (Matsuoka et al., 2007). The PIKK-family kinases have both redundant and distinct functions. ATM and DNA-PK function in response to DSBs throughout the cell cycle, and ATR functions mostly in DNA replication-associated damage during the S phase (Gospodinov and Ugrinova, 2019; Riabinska et al., 2013). Even though these kinases all prefer to phosphorylate a serine or threonine residue that is followed by a glutamine (SQ/TQ motif) (Blackford and Jackson, 2017), their knockout phenotypes are different. ATM-knockout mice are sterile and often suffer from lymphopenia, whereas DNA-PK knockout mice are fertile and have the severe combined immunodeficiency (SCID) phenotype (Menolfi and Zha, 2020). It has been proposed that ATM promotes γ-H2AX clustering and DNA repair accuracy, whereas DNA-PK is essential for end joining (Caron et al., 2015). ATM is also known to phosphorylate threonine 392 of males absent on the first (MOF) (Gupta et al., 2014) that is a histone acetyltransferase for H4 lysine 16 acetylation (H4K16ac) (Sharma et al., 2010; Taipale et al., 2005), which can assist in chromatin decompaction and facilitate recruitment of DNA repair machinery, including homologous recombination (HR) repair proteins (Dhar et al., 2017; Gupta et al., 2014; Horikoshi et al., 2019; Hunt et al., 2013; Kim et al., 2019; Sharma et al., 2010). However, it remains unclear how ATM and DNA-PK function in γ-H2AX formation just after DSBs are generated.

The dynamics of γ-H2AX have been analyzed by immunofluorescence, immunoblotting, and chromatin immunoprecipitation (Burma et al., 2001; Caron et al., 2015; Stiff et al., 2004), but the γ-H2AX-formation kinetics immediately after DSBs (within a few minutes) in single cells have not been elucidated because of the lack of a monitoring system for γ-H2AX in living cells. However, by introducing fluorescently labeled modification-specific antigen-binding fragments (Fabs) into living cells, changes in the target modifications can be tracked (Conic et al., 2018; Hayashi–Takanaka et al., 2011; Sato et al., 2021). In the present study, we used γ-H2AX-specific Fabs (Yamagata et al., 2019) to analyze the kinetics of γ-H2AX formation in response to laser-induced DNA damage in living human cells. We first compared the involvement of ATM in immediate γ-H2AX kinetics. After laser microirradiation, γ-H2AX accumulated at the damaged areas in similar kinetics in ATM-proficient or -deficient cells. In contrast, the inhibition of DNA-PK activity slowed the γ-H2AX accumulation and dissolution kinetics. Fluorescence recovery after photobleaching (FRAP) and permeabilized cell assays revealed that a DNA-PK subunit diffused freely throughout the nucleus, whereas ATM repeatedly binds to and unbinds from chromatin. Thus, DNA-PK can bind DNA ends immediately after DSBs and phosphorylate H2AX whereas ATM appears to be dispensable for immediate early γ-H2AX accumulation in DSBs.

## RESULTS

### Kinetics of γ-H2AX immediately after DSB induction

To examine the immediate early kinetics of γ-H2AX in response to DNA damage, we loaded fluorescently labeled γ-H2AX-specific Fabs that had been used to detect DSBs in mouse embryos (Yamagata et al., 2019) into cells and microirradiated a part of the nucleus using a 405-nm laser to induce the DSBs (Muster et al., 2017) (Fig. 1A). We first compared the kinetics in ATM-deficient (AT5BIVA) and -proficient (11-4) cells. AT5BIVA cells harbor an in-frame deletion in the kinase domain of the ATM gene (Cheema et al., 2013; Gilad et al., 1996), resulting in little ATM protein expression (Fig. S1). The 11-4 cells were derived from AT5BIVA by transferring chromosome 11 to express the functional ATM (Komatsu et al., 1990) (Fig. S1). Before laser microirradiation, Cy5-labeled γ-H2AX Fabs were distributed throughout the cytoplasm and nucleus except for the nucleoli in both AT5BIVA and 11-4 cells (Fig. 1B, −0:15 min:sec). Just after 405-nm laser irradiation, the fluorescence in the irradiated area was decreased by photobleaching (0:00) and then increased over the nuclear background within minutes (0:15, 4:00, and 7:30). The relative fluorescence intensities of γ-H2AX Fab in the irradiated area were measured and plotted. In both ATM-deficient (AT5BIVA) and - proficient (11-4) cells, γ-H2AX Fabs accumulated in the irradiated area reaching a broad peak at around 100–200 s and then gradually declined (Fig. 1C). The γ-H2AX Fabs showed slightly higher accumulation in the 11-4 cells than in the AT5BIVA cells, but not significantly (*p* = 0.0881 at time 180 s).

**Fig. 1.**
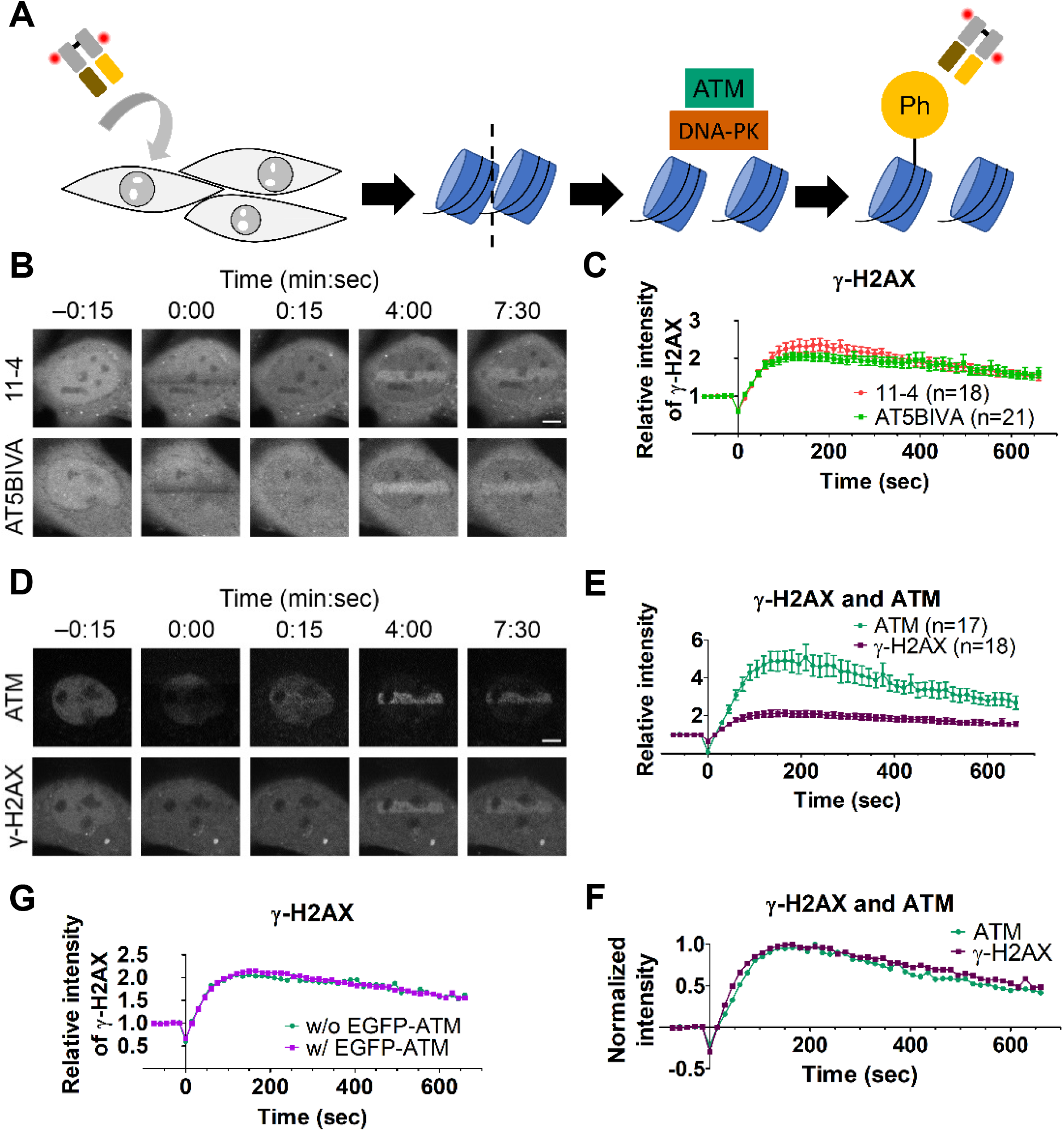
Dynamics of ATM and γ-H2AX after DSB induction by 405-nm laser bleaching. (A) Schematic diagram of the experimental system showing how phosphorylated H2AX (γ-H2AX) in response to a DSB is detected in living cells. A fluorescent dye-conjugated γ-H2AX-specific Fab is loaded into cells, and then DNA damage is induced by 405-nm laser irradiation. ATM and DNA-PK are recruited to DNA lesions and phosphorylate histone H2AX to form γ-H2AX, which is now recognized by the Fab. Thus, the localization and degree of γ-H2AX can be monitored by live-cell imaging. (B and C) Monitoring of γ-H2AX in response to laser microirradiation in ATM-proficient (11-4) and -deficient (AT5BIVA) cells. Cy5-conjugated γ-H2AX Fab had been loaded into 11-4 and AT5BIVA cells and rectangular areas were then irradiated to induce DSBs. (B) Time-lapse images. In the irradiated areas, fluorescence was bleached just after irradiation (0:00) and then increased (4:00 and 7:30). (C) Accumulation kinetics of γ-H2AX. The relative intensities of γ-H2AX Fab in the irradiated areas in 11-4 and AT5BIVA cells are plotted (mean ± SEM, with the number of cells, n). (D–F) Monitoring EGFP-ATM and γ-H2AX in response to laser microirradiation. (D) Time-lapse images. AT5BIVA cells that express EGFP-ATM plasmid were loaded with Cy5-conjugated γ-H2AX Fab and rectangular areas were irradiated to induce DSBs. Both EGFP-ATM and γ-H2AX accumulated in the irradiated areas. Scale bar: 5 μm. (E and F) Accumulation kinetics of EGFP-ATM and γ-H2AX. The relative intensities (mean ± SEM, with the number of cells, n) (E) and normalized intensities from the baseline before irradiation to the maximum (F) in the irradiated area are plotted. (G) Comparison of the accumulation kinetics of Cy5-conjugated γ-H2AX Fab in irradiated areas in AT5BIVA cells without and with EGFP-ATM, reproduced from data in (C) and (E). No difference is seen. Scale bars: 5 μm.

To compare the accumulation kinetics of ATM and γ-H2AX in damaged areas, EGFP-tagged ATM (EGFP-ATM) was expressed in AT5BIVA cells. Accumulation in the irradiated area was greater for EGFP-ATM than for γ-H2AX (Figs. 1D,E), but when normalized by setting the maximum intensity at 1 and the original intensity at 0, the accumulation kinetics of EGFP-ATM and γ-H2AX were similar and ATM accumulation was slightly delayed relative to that of γ-H2AX (Fig. 1F). The kinetics of γ-H2AX did not change in the AT5BIVA cells without or with EGFP-ATM expression (Fig. 1G). These data suggest that ATM is not essential for H2AX phosphorylation immediately after DSBs were induced by laser irradiation, although the accumulation of ATM coincided with the γ-H2AX kinetics.

We next determined if the kinetics of γ-H2AX accumulation differ in different cell-cycle phases because ATR is known to function during the S phase. In addition, the accumulation kinetics could be affected by the preference of DSB repair pathways, HR or non-homologous end joining (NHEJ), which depends on the cell cycle (Her and Bunting, 2018; Karanam et al., 2012; Shrivastav et al., 2008). For a cell-cycle marker in living cells, we used PCNA tagged with a fluorescent protein, which shows characteristic distributions depending on the cell-cycle phase (Essers et al., 2005; Leonhardt et al., 2000; Mir et al., 2011; Schonenberger et al., 2015; Somanathan et al., 2001). mCherry-PCNA diffused in both the cytoplasm and nucleus during most of the G1 phase, concentrated on foci during the S phase, and diffused in the nucleus during the G2 phase (Figs. S2A–B) (Uchino et al., 2022). The accumulation kinetics of γ-H2AX Fabs were essentially similar in all cell-cycle phases (Figs. S2C–G), with slightly more accumulation in 11-4 cells than AT5BIVA cells as observed without mCherry-PCNA expression (Fig. 1C).

### Effects of the specific inhibitors of ATM, ATR, and DNA-PK on H2AX phosphorylation

The finding that similar γ-H2AX accumulation kinetics were observed with and without ATM suggested that other PIKK-family kinases (ATR and/or DNA-PK) might have a primary role in immediate H2AX phosphorylation or compensate for ATM function in deficient cells. To determine which of these PIKK-family kinases is a primary kinase for immediate γ-H2AX formation, we first determined the effective concentration of kinase-selective inhibitors in AT5BIVA and 11-4 cells by immunofluorescence. The cells were incubated with an ATR inhibitor (AZ20), an ATM inhibitor (KU55933), or a DNA-PK inhibitor (NU7441) at 10, 5, 2.5, and 0 μM, simultaneously with 20 μg/mL etoposide (ETP) to induce DSBs for 1 h, before fixing and staining with γ-H2AX-specific antibody (Figs. S3A–C). In ETP-treated 11-4 cells, γ-H2AX signals were observed at similar levels in the presence of a single inhibitor (Fig. S3D). In AT5BIVA cells, γ-H2AX fluorescence intensity was drastically decreased in the DNA-PK inhibitors, but not in ATR and ATM inhibitors (Fig. S3E). Susceptibility to the DNA-PK inhibitor in AT5BIVA cells was rescued by EGFP-ATM expression (Fig. S3F). This result is consistent with a previous study showing that in the absence of functional ATM, γ-H2AX formation after DSBs was primarily mediated through DNA-PK (Stiff et al., 2004). When DNA-PK activity is inhibited, ATM can phosphorylate H2AX in 60 min.

To analyze the effect of DNA-PK on immediate early γ-H2AX formation kinetics, cells were incubated with 2.5 μM NU7441 for ≥1 h before and during the laser-irradiation assay. To minimize the contribution of ATR, we chose G1 cells for the analysis using cells stably expressing mCherry-PCNA. In 11-4 cells treated with NU7441, γ-H2AX Fab continually accumulated for ∼500 s, in contrast to the decrease after ∼200 s in control cells without the inhibitor (Figs. 2A,B). The delayed decrease may have been caused by ATM hyperactivation that can be induced by DNA-PK inhibitors (Zhou et al., 2017) or by impaired DNA repair. In AT5BIVA cells treated with NU7441, in which both ATM and DNA-PK activities are diminished, γ-H2AX Fab accumulation was substantially retarded compared to that in control ATM-deficient cells (Figs. 2A,C). This result indicated that DNA-PK activity is critical in the early response to DSBs. The basal activity of ATR (Boe et al., 2018) might mediate the low level of γ-H2AX accumulation in NU7441-treated AT5BIVA cells (also observed in immunofluorescence in Fig. S3C).

**Fig. 2.**
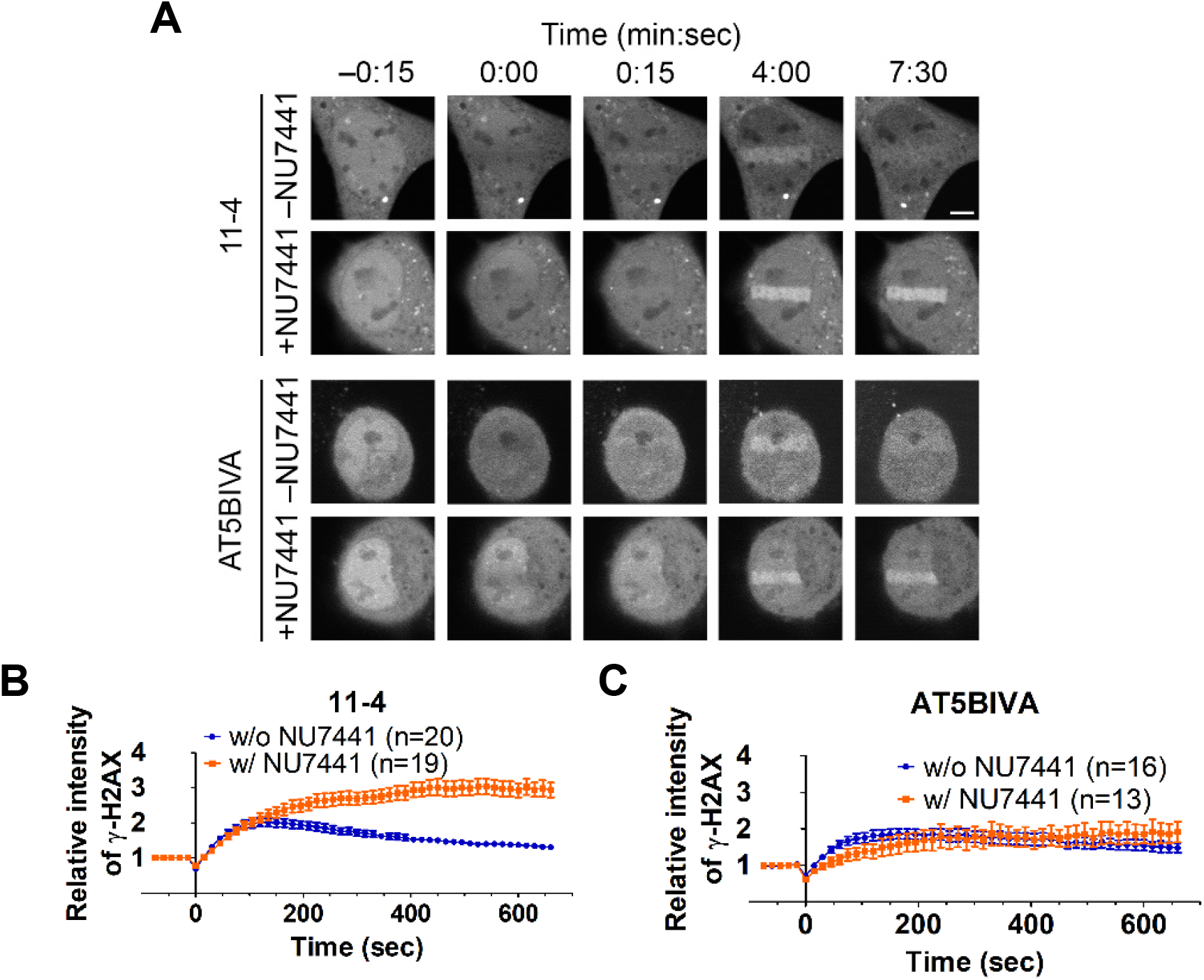
Effect of DNA-PK inhibitor on γ-H2AX dynamics. Loading of 11-4 and AT5BIVA cells with Cy5-conjugated γ-H2AX Fab that were then treated with or without 2.5-μM NU7441 for ≥1 h before and during the laser-irradiation assay. (A) Time-lapse images. (B and C) Accumulation kinetics of γ-H2AX. The relative intensities of γ-H2AX Fab at the irradiated areas in 11-4 (B) and AT5BIVA (C) are plotted (mean ± SEM, with the number of cells, n). Scale bars: 5 μm.

### Involvement of MOF in early DNA damage response

Crosstalk between γ-H2AX and another histone modification, H4K16ac, has been previously demonstrated. H4K16ac is mediated through a histone acetyltransferase, MOF (Sharma et al., 2010; Taipale et al., 2005), and is involved in DSB repair (Dhar et al., 2017; Horikoshi et al., 2019; Kim et al., 2019; Miller et al., 2010). H4K16ac obstructs the binding of 53BP1 to H4K20me2 (Tang et al., 2013) and regulates the DNA repair pathway choice by limiting the DNA end resection which is a required step for HR (Pellegrino et al., 2017). ATM phosphorylates MOF to facilitate HR protein recruitment (Gupta et al., 2014) and MOF facilitates ATM kinase activity (Gupta et al., 2005; Li et al., 2010). Therefore, we analyzed the possible involvement of H4K16ac in early γ-H2AX kinetics. We first used the specific Fab to determine if H4K16ac is accumulated in laser-irradiated areas. After the induction of DSBs, H4K16ac Fab did not show obvious accumulation in either AT5BIVA or 11-4 cells regardless of the presence of the DNA-PK inhibitor, NU7441 (Figs. S4A,B). This suggests that H4K16ac levels do not change several min after DSB induction.

We next used a lentivirus-mediated shRNA expression system to investigate the function of MOF in ATM- and/or DNA-PK-mediated H2AX phosphorylation, by knocking down MOF in AT5BIVA, 11-4, and AT5BIVA expressing EGFP-ATM. Given that MOF knockdown increased apoptotic cells (Li et al., 2012; Thomas et al., 2008), we analyzed cells that showed normal nuclear shape regardless of the cell-cycle phase. Immunoblotting showed that MOF-specific shRNA expression lowered MOF and H4K16ac levels to 10%–20% and 20%–40%, respectively, relative to the expression of scramble shRNA control (Figs. 3A–C). Immunofluorescence confirmed the decrease in H4K16ac by MOF-specific shRNA expression (Fig. 3D and Fig. S5) and showed that γ-H2AX was still formed by ETP treatment for 20 min in MOF knockdown cells (Fig. S5A–C). In a laser-irradiation assay,γ-H2AX Fab kinetics in 11-4 cells were similar in the MOF knockdown and the scrambled shRNA control cells (Fig. 3E), suggesting that MOF did not affect the accumulation kinetics of γ-H2AX when both ATM and DNA-PK were present, in agreement with a previous report (Li et al., 2010). When DNA-PK was inhibited by NU7441 in 11-4 cells, γ-H2AX accumulation was also similar in MOF knockdown and control cells, although a subtle delay in MOF knockdown was observed (Fig. 3F), suggesting that inhibition of DNA-PK has a greater effect than MOF knockdown. In AT5BIVA cells, γ-H2AX accumulation was not affected by MOF knockdown (Fig. 3G). When AT5BIVA cells were treated with a DNA-PK inhibitor, combined with MOF knockdown, γ-H2AX barely accumulated (Fig. 3H). Since ATR activity remained in AT5BIVA cells treated with the DNA-PK inhibitor, the repression of H2AX phosphorylation by MOF knockdown suggests that ATR activity might also be regulated by MOF.

**Fig. 3.**
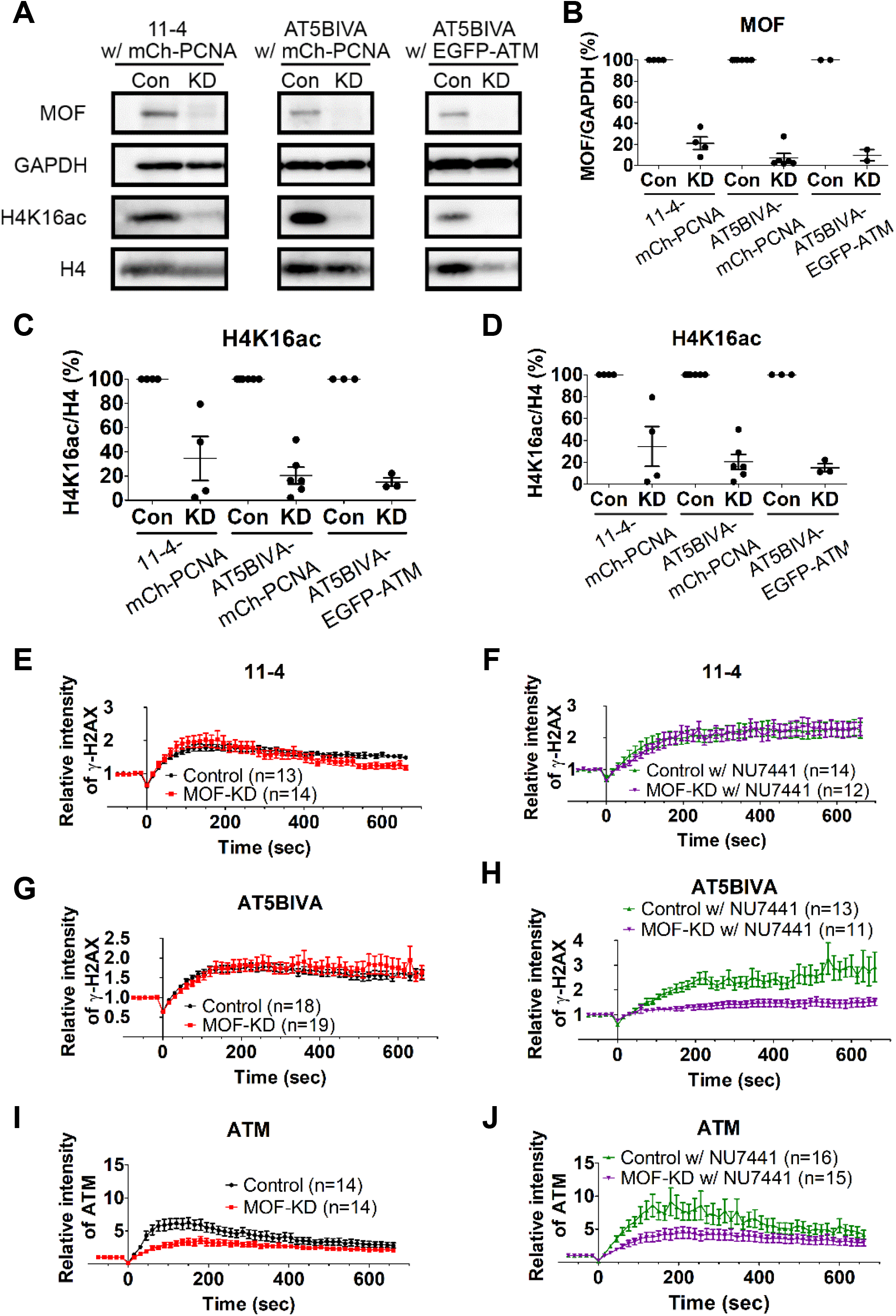
Effects of MOF knockdown on the accumulation of γ-H2AX. Lentivirus was used to infect 11-4 and AT5BIVA cells that express mCherry-PCNA or AT5BIVA cells that express EGFP-ATM to cause expression of MOF-specific and control-scramble shRNA. (A–D) Validation of MOF knockdown. (A) Immunoblots showing decreased levels of MOF. Con, control-scramble shRNA expression; KD, MOF-knockdown. (B and C) Relative amounts of MOF and H4K16ac evaluated by immunoblotting. The band intensities of MOF (B) and H4K16ac (C) are normalized against those of GAPDH and histone H4, respectively, and further normalized using Con. Plots showing four biologically independent experiments (mean ± SEM, with individual data points). (D) Fluorescence intensity of H4K16ac in MOF knockdown (mean ± SEM; n = 3 independent experiments indicated by different colors; ≥115 cells were analyzed in each experiment). (E–H) Accumulation kinetics of γ-H2AX. (E) 11-4 cells with MOF-KD and Con. (F) 11-4 cells in 2.5 μM NU7441 with MOF-KD and Con. (G) AT5BIVA cells with MOF-KD and Con. (H) AT5BIVA cells in 2.5 μM NU7441 with MOF-KD and Con. (I and J) Accumulation kinetics of ATM in AT5BIVA cells that express EGFP-ATM. (I) Cells without NU7441 with MOF-KD and Con. (J) Cells in 2.5 μM NU7441 with MOF-KD and Con. Means ± SEM with the number of cells (n) are shown.

The effects of MOF knockdown on ATM accumulation kinetics in laser-irradiated areas were also analyzed using AT5BIVA cells expressing EGFP-ATM (Figs. 3I–J). The accumulation of EGFP-ATM in the irradiated area was greater in MOF-knockdown cells than in the scrambled control without and with NU7441 (Figs. 3I–J). This result supports the role of MOF as a regulator of ATM (Gupta et al., 2005).

### Mobility of ATM and Ku80, a subunit of DNA-PK, in living cells

We next investigated the mobility of ATM and Ku80, a subunit of DNA-PK, and their response to DNA damage, using FRAP with a 488-nm laser, which does not induce DSBs as does a 405-nm laser (Arimura et al., 2013; Muster et al., 2017). Without DNA damage, EGFP-ATM recovered in a few seconds after bleaching. EGFP-Ku80 recovered within 0.5 s, much faster than EGFP-ATM (Figs. 4A,B). In the area with DSBs that were induced by 405-nm laser irradiation, EGFP-ATM recovery became much slower, whereas the recovery kinetics of EGFP-Ku80 remained unchanged (Figs. 4A,C). These results suggest that EGFP-ATM repeatedly binds to and dissociates from chromatin and when DSBs are induced, EGFP-ATM binds more stably to chromatin. In contrast, EGFP-Ku80 appears to diffuse almost freely in the nucleus. The finding of little or no change in EGFP-Ku80 kinetics in DSBs can be explained by the transient binding of DNA-PK to damaged chromatin and/or by a tiny fraction of DNA-PK bound to damaged chromatin. In fact, EGFP-Ku80 was not enriched in irradiated areas under the conditions used (see below for the results with more damage).

**Fig. 4.**
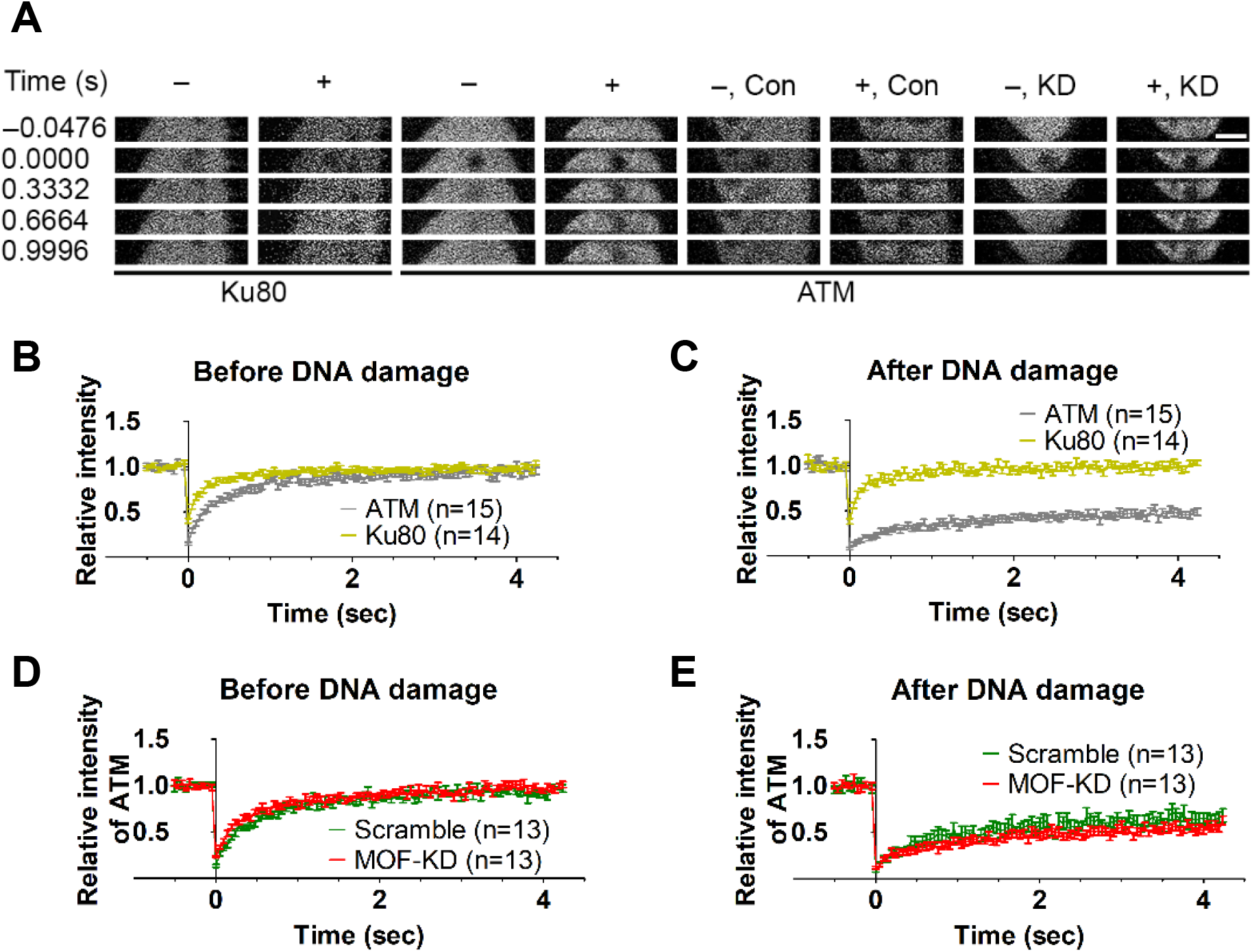
MOF does not affect ATM binding to the DNA-damaged region. FRAP using a 488-nm laser was performed for EGFP-ATM in AT5BIVA and EGFP-Ku80 in 11-4 cells. (A) Example images. –, without DNA damage induction; +, with DNA damage induction by 405-nm laser irradiation; Con, control-scramble shRNA expression; KD, MOF-knockdown. (B–E) FRAP recovery curves (mean ± SEM) without (B and D) and with laser irradiation (C and E). (B) EGFP-ATM and EGFP-Ku80. (C) EGFP-ATM and EGFP-Ku80 in irradiated areas. (D) EGFP-ATM with MOF-KD and Con. (E) EGFP-ATM in irradiated areas with MOF-KD and Con. Means ± SEM with the number of cells (n) are shown. Scale bars: 5 μm.

We also investigated whether or not MOF knockdown affects ATM mobility (Figs. 4A,D–E). MOF knockdown had little effect on the recovery kinetics without or with DNA damage (Figs. 4A,D–E). Thus, the residence time of EGFP-ATM on both undamaged and damaged chromatin does not depend on MOF and H4K16ac, whereas the kinase activity of ATM at damaged chromatin appears to be facilitated by MOF (Fig. 3I).

### ATM, but not DNA-PK, was observed to bind to chromatin in permeabilized cells

To further analyze the different dynamics of ATM and DNA-PK and the relevance to H2AX phosphorylation, we used a permeabilized cell system. When cells are permeabilized with a non-ionic detergent, such as TritonX-100, freely diffusible proteins are extracted, whereas chromatin-bound proteins remain (Jackson and Cook, 1985; Kimura et al., 2006; Nickerson et al., 1997). Permeabilized 11-4 and AT5BIVA cells were incubated with γ-H2AX Fab, and DNA damage was then induced by laser irradiation (Fig. 5A). Accumulation of γ-H2AX Fab in the damaged area was observed in permeabilized 11-4 cells although it was much slower than that in intact cells (Fig. 5B). By contrast, γ-H2AX Fab did not accumulate in permeabilized AT5BIVA cells (Fig. 5B). These results are consistent with the view that ATM transiently binds to chromatin so that a chromatin-bound fraction remains during permeabilization, whereas DNA-PK freely diffuses without DNA damage and is mostly extracted during permeabilization (Fig. 5A). Thus, in permeabilized cells, only ATM-proficient cells contain H2AX phosphorylation activity in response to laser-induced DNA damage (Fig. 5A). Under MOF knockdown, the accumulation of γ-H2AX Fab in 11-4 cells was impaired (Fig. 5C), suggesting that chromatin-bound MOF and/or H4K16ac is required for ATM-mediated H2AX phosphorylation. MOF knockdown did not affect γ-H2AX Fab accumulation in AT5BIVA cells (Fig. 5D), which is reasonable because ATM is lacking.

**Fig. 5.**
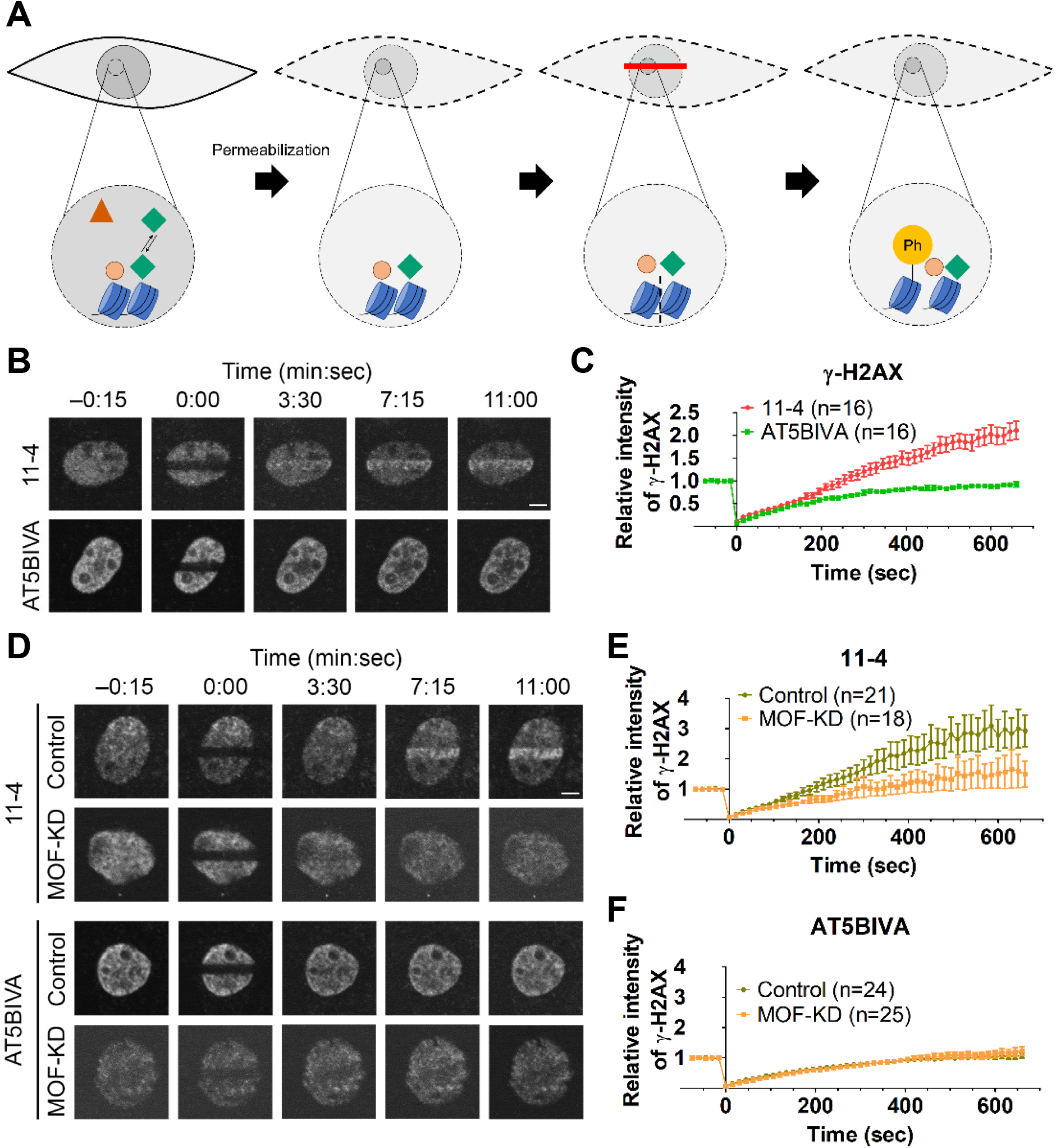
H2AX phosphorylation activity in permeabilized cells. (A) Schematic diagram of cell permeabilization and H2AX phosphorylation. Within the intact cells, some proteins (indicated in the red triangle) diffuse freely and others transiently bind to the chromatin (the first panel). When cells are permeabilized with 0.1% TritonX-100, only chromatin-unbound proteins are extracted (second panel). After DSBs are induced by laser irradiation, histone H2AX becomes phosphorylated if a kinase remains on chromatin in permeabilized cells (third and fourth panels). The γ-H2AX in permeabilized cells can be detected by the accumulation of dye-labeled specific Fab. (B and C) 11-4 and AT5BIVA cells were permeabilized and laser-irradiated in the presence of Cy5-conjugated γ-H2AX Fab. Time-lapse images (B) and accumulation kinetics of γ-H2AX at the irradiated areas (C) are shown. (D-F) MOF was knocked down in 11-4 and AT5BIVA cells before permeabilization and laser irradiation. (D) Time-lapse images. (E-F) Accumulation of γ-H2AX at the irradiated areas in 11-4 (E) and AT5BIVA (F). Means ± SEM with the number of cells (n) are shown. Scale bars: 5 μm.

### Effects of massive DSBs induced by laser irradiation in sensitized cells

Finally, we examined the kinetics of γ-H2AX, EGFP-ATM, and EGFP-Ku80 using Hoechst-sensitized cells in which enhanced DNA damage can be induced by laser irradiation (Bekker– Jensen et al., 2006). Within several seconds, γ-H2AX was accumulated in irradiated areas in sensitized 11-4 cells and was then soon enriched in unirradiated areas in the nucleus (Fig. S6A). Unlike previous conditions in unsensitized cells, EGFP-Ku80 accumulated in laser-irradiated areas, and its accumulation was more rapid than accumulation of γ-H2AX in 11-4 cells and EGFP-ATM in AT5BIVA cells (Figs. 6A–B, Fig. S6B). These data are consistent with the view that abundant and diffused Ku80 rapidly recognizes broken DNA ends and that DNA-PK initiates H2AX phosphorylation (Davis et al., 2010; Kochan et al., 2017). Treatment with the DNA-PK inhibitor, NU7441, did not affect the dynamics of EGFP-Ku80 (Figs. 6C–E), suggesting that Ku80 accumulation in the damaged region does not depend on the kinase activity of DNA-PK. The mobility of EGFP-Ku80 determined by FRAP in sensitized cells became slightly slower after laser irradiation, suggesting that the residence time on damaged chromatin is shorter for DNA-PK than for ATM, which is in good agreement with a previous report (Davis et al., 2010).

**Fig. 6.**
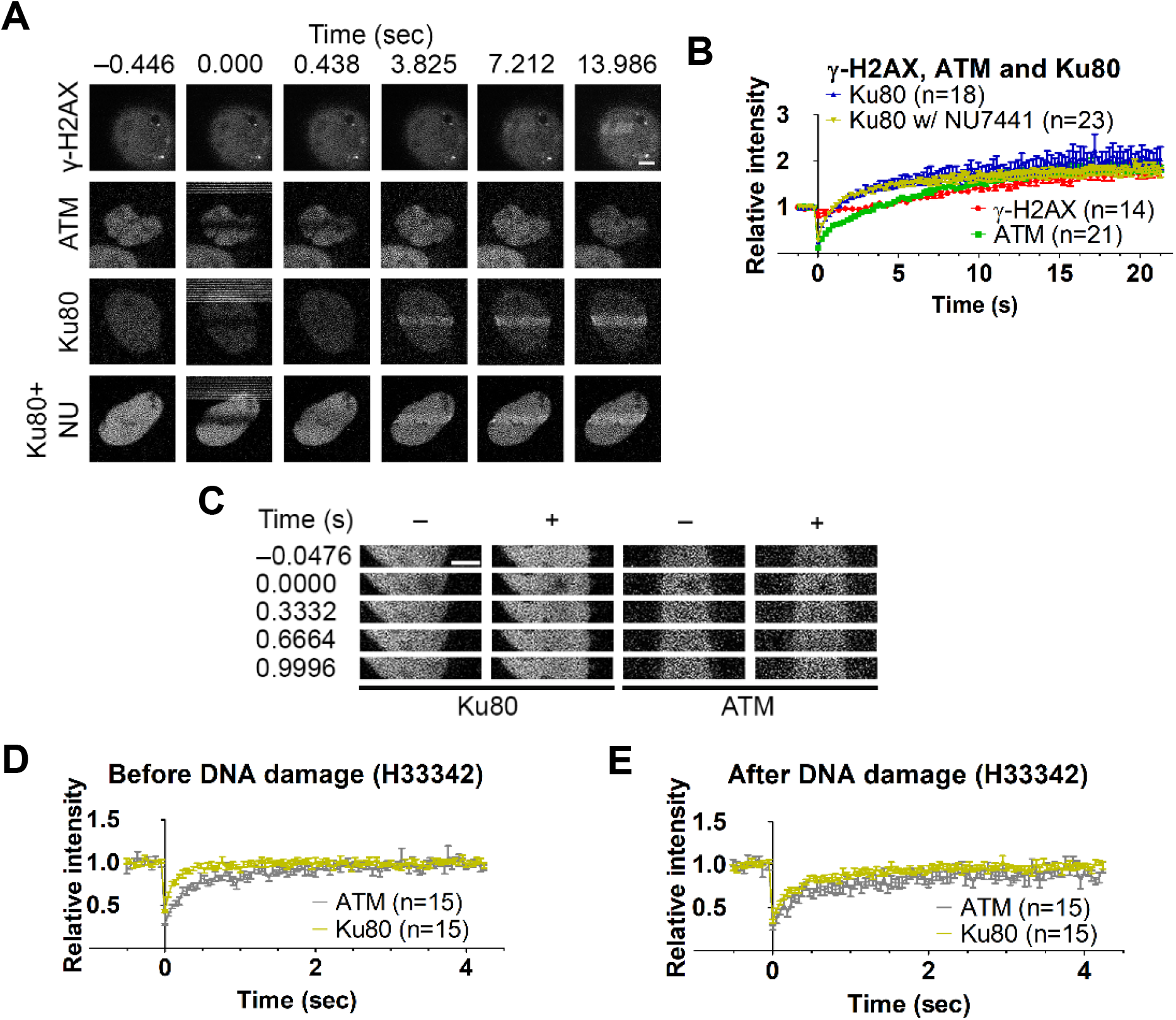
Effects of massive DSBs induced by Hoechst 33342 sensitization following laser irradiation. DSBs were induced by irradiating Hoechst 33342-sensitized cells. Time-lapse images were acquired for γ-H2AX Fab and EGFP-Ku80 (± 2.5 μM NU7441) in 11-4 cells, and EGFP-ATM in AT5BIVA cells. (B) Accumulation of γ-H2AX, Ku80, and Ku80 in 2.5-μM NU7441, and of EGFP-ATM in irradiated areas in Hoechst 333420-sensitized cells (mean ± SEM). (C–E) FRAP with a 488-nm laser without and with DSB. (C) Time-lapse images of EGFP-Ku80 and EGFP-ATM before and after bleaching. (D and E) Fluorescence recovery without (D) or with DSBs (E). Means ± SEM with the number of cells (n) are shown. Scale bars: 5 μm.

**Fig. 7.**
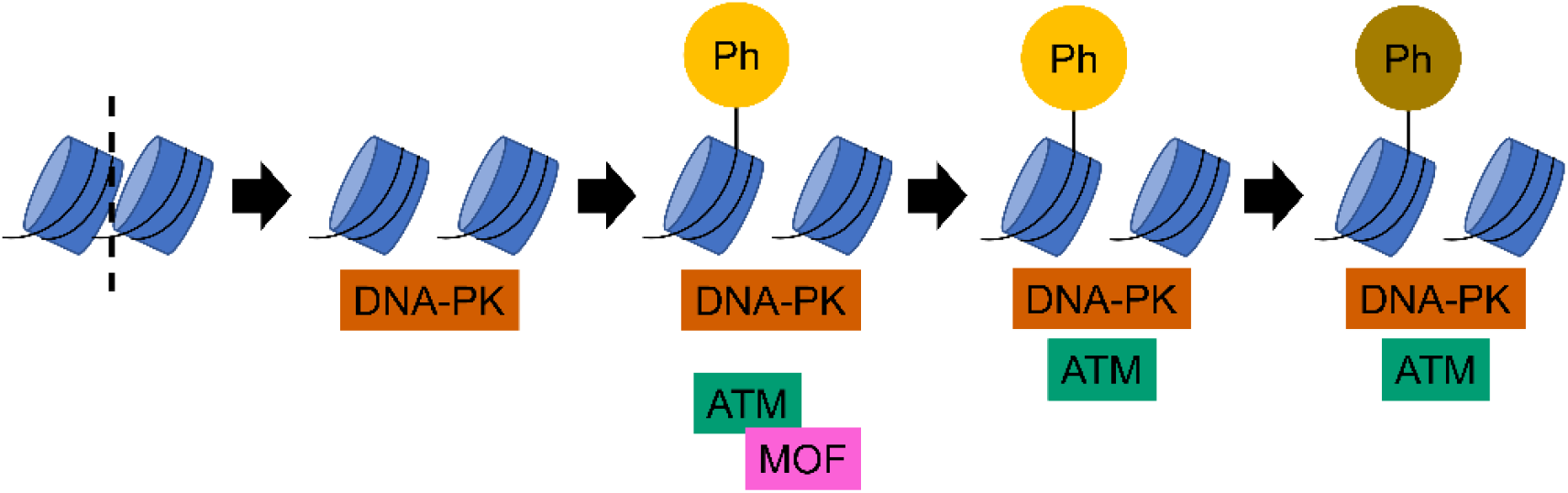
Model of early γ-H2AX dynamics in response to DSB. After DSB, DNA-PK that diffuses throughout the nucleus binds to DNA ends and phosphorylates histone H2AX. ATM also phosphorylates H2AX and other proteins assisted by MOF, although accumulation at damaged sites is slower for ATM than for DNA-PK.

## DISCUSSION

ATM is known as a major kinase that can phosphorylate the histone H2A variant, H2AX, to give γ-H2AX in response to DSB (Burma et al., 2001; Caron et al., 2015), but the mechanism by which γ-H2AX is formed immediately after DSB is not fully understood. In this study, we employed Fab-based endogenous modification labeling (Hayashi–Takanaka et al., 2011) to detect the rapid formation of γ-H2AX in response to DNA damage induced by laser irradiation. In both ATM-proficient and -deficient cells, γ-H2AX accumulated immediately after irradiation and reached a broad peak at ∼100–200 s before gradually decreasing, which may be associated with progression of DNA repair (Bouquet et al., 2006; MacPhail et al., 2003; Mah et al., 2010). Thus, ATM does not appear to have a major role in the immediate γ-H2AX formation upon DNA damage, although ATM has a critical role in later responses (Caron et al., 2015; Kuhne et al., 2004; Lobrich and Jeggo, 2005; Loucas and Cornforth, 2004; Riballo et al., 2004; Stiff et al., 2004). ATM can amplify the DNA damage signal that is initially generated via DNA-PK (Lu et al., 2019) and may be critical for heterochromatin repair that requires a longer time (Goodarzi et al., 2008). Inhibition of DNA-PK using NU7441 altered the kinetics of γ-H2AX to continually accumulate for ≤400–500 s in ATM-proficient cells, but accumulation was drastically delayed in ATM-deficient cells. These results support the critical role of DNA-PK in the immediate DNA damage response (Caron et al., 2015; Lu et al., 2019; Riballo et al., 2004), including γ-H2AX phosphorylation (Liu et al., 2019). DNA-PK has also been shown to function in chromatin decompaction and initiation of the DSB response (Lu et al., 2019). Here, we demonstrated that the Ku80 subunit of DNA-PK that diffuses freely in the nucleus. It is likely that as soon as DSBs are induced, DNA-PK binds to DNA ends and phosphorylates H2AX more rapidly than ATM, which transiently binds to chromatin in the steady state (Aleksandrov et al., 2018; Davis et al., 2010; Kochan et al., 2017). Once γ-H2AX is formed, it facilitates the binding of ATM to phosphorylate nearby H2AX and DNA repair proteins such as NBS1 (Lim et al., 2000) and p53 (Banin et al., 1998). In the absence of DNA-PK activity, γ-H2AX is still formed via ATM that can survey chromatin by repeated association and dissociation and can also bind to DNA repair machinery (Callen et al., 2009). Furthermore, ATM becomes hyperactivated by DNA-PK inhibition (Finzel et al., 2016). The slowed and prolonged γ-H2AX accumulation in cells in which DNA-PK activity is inhibited might be caused by this ATM hyperactivation, which later affects the downstream response of the DNA damage pathway, including the p53 pulse (Finzel et al., 2016; Sun et al., 2017). Although chromatin binding of ATM does not depend on MOF, immediate activation of ATM and ATR does require MOF (Gupta et al., 2005) and H4K16ac may function in later stages of DNA repair (Miller et al., 2010). Thus, the distinct binding and activation mechanisms of ATM and DNA-PK may contribute to the DSBs response through overlapping functions in H2AX phosphorylation.

The DSB repair pathway choice depends on the cell cycle (Her and Bunting, 2018; Karanam et al., 2012). NHEJ occurs throughout the cell cycle but preferentially functions during the G1 phase (Lieber et al., 2003; Mao et al., 2008b), whereas HR that uses the sister chromatid as a template for repair can be used in the S and G2 phases (Kadyk and Hartwell, 1992). Although the difference in the DNA repair pathway might affect the dynamics of γ-H2AX (Shrivastav et al., 2008), the immediate early accumulation kinetics were similar in all cell-cycle phases with or without ATM. Taken together with the critical role of DNA-PK, this observation is consistent with the idea that the preferred pathway in mammalian cells is NHEJ for faster and more efficient repair processes (Mao et al., 2008a; Mao et al., 2008b).

Our results show that using γ-H2AX-specific Fabs is a powerful tool to study the early dynamics of DSB. However, since using the ATM inhibitor does not phenocopy the ATM-mutated cells (Choi et al., 2010), the DNA-PK inhibitor treatment might also not phenocopy the DNA-PK-deficient cells. Therefore, the future study should investigate whether the delay in the accumulation of γ-H2AX is still observed in the DNA-PK-deficient cells or not. In addition, since the dynamics of p53 differs in different cell lines (Stewart–Ornstein and Lahav, 2017), the future study should address whether the γ-H2AX dynamics depends on p53. In summary, our results confirm that ATM is dispensable for the immediate early DSB, and support the significance of DNA-PK in the early response after DSB is induced.

## MATERIALS AND METHODS

### Cell culture

AT5BIVA, an SV40-transformed AT fibroblast cell line, and 11-4 cells, which are AT5BIVA cells with the addition of chromosome 11 to restore ATM, were gifts from S. Tashiro (Hiroshima University) (Sun et al., 2010). HEK293T was obtained from K. Fujinaga (Sapporo Medical University). AT5BIVA, 11-4, and HEK293T cells were routinely cultured in Dulbecco’s Modified Eagle Medium (DMEM; Nacalai Tesque) supplemented with 10% fetal bovine serum (FBS; Thermo Fisher Scientific) and 1% L-Glutamine-Penicillin-Streptomycin solution (GPS; Sigma-Aldrich) at 37°C under 5% CO2.

### Plasmids and transfection

Cells were plated in a 6-well plate (Thermo Fisher Scientific) one day before the transfection of the expression construct of mCherry-proliferating-cell nuclear antigen (PCNA) or EGFP-ATM based on PB533A-2 (System Biosciences) with a PiggyBac transposon expression vector (PB210PA-1; System Biosciences), or pEGFP-C1-FLAG-Ku80 ((Britton et al., 2013), Addgene; 46958) using FuGENE HD Transfection Reagent (Promega) according to the manufacturer’s instruction. The EGFP-ATM expression vector was constructed using a plasmid containing ATM cDNA provided by T. Ikura (Kyoto University) and the entire ATM sequence was verified by sequencing. To obtain a stable cell line, 2 days after the transfection, the cells were incubated in the presence of 1 mg/mL G418 Disulfate Aqueous Solution (Nacalai Tesque) in the complete medium for >1 week. Cells that exhibited mCherry-PCNA or EGFP-ATM fluorescence were sorted using a cell sorter (Sony; SH800) and cultured in a fresh medium without G418.

### Lentiviral shRNA infection

HEK293T cells were plated 1 day before transfection with psPax2 (viral packaging plasmid; Addgene; 12260), pCMV-VSV-G (viral envelope plasmid; Addgene; 8454), and a pLKO.1-based plasmid containing either scrambled sequences or hMOF shRNA (Kapoor–Vazirani et al., 2008; Kapoor–Vazirani et al., 2011) (pCMV-VSV-G and pLKO.1 puromycin were gifts from R. Weinberg through Addgene), using Lipofectamine 3000 (Invitrogen) according to the manufacturer’s instruction. The pLKO.1-based plasmids were constructed according to the Addgene’s protocol. Medium containing lentiviral particles collected 1 day after transfection were filtered through a 0.45-μm filter (Advantec), and then 2 μg/mL polybrene (Sigma-Aldrich) was added. Recipient cells were plated one day before the infection and the medium was replaced with a lentiviral-containing medium for 18–24 hours. Then, the medium was changed to a fresh medium containing 1 μg/mL puromycin (Thermo Fisher Scientific) for 11-4 cells and 2 μg/mL puromycin for AT5BIVA cells. After 1–2 days of puromycin selection, the medium was replaced with a fresh medium.

### Live-cell imaging

Cells were plated on a 35-mm glass-bottom dish with a coverslip (AGC Techno Glass). The next day, fluorescent dye-labeled modification-specific Fab was loaded into cells using glass beads (Hayashi–Takanaka et al., 2011) and the medium was changed to FluoroBrite DMEM (Thermo Fisher Scientific) supplemented with 10% FBS and 1% GPS. The preparation and dye-conjugation of Fabs have been described previously (Kimura and Yamagata, 2015; Yamagata et al., 2019). A glass-bottom dish was set on a heated stage (Tokai Hit) with a CO2 control system (Token) on a confocal microscope (FV-1000, Olympus) operated by built-in software (Fluoview ver. 4.2) with a PlanSApo 60× (NA 1.40) oil-immersion objective lens to maintain cells at 37°C under 5% CO2. For the laser-irradiation assay, five images were collected using the line-sequential imaging mode (512 × 512 pixels; pinhole 800 μm; 8× zoom; 2-line Kalman filtration) with three laser lines (0.1%–2.0% 488-nm laser transmission; 0.5%– 10.0% 543-nm laser transmission; 0.5%–5.0% 633-nm laser transmission), then a 26.06 × 2.25-μm rectangle area was irradiated with 100% 405-nm laser transmission for 3.09 sec, and another 45 images were collected using the original settings at 15-sec intervals. The time-series images were aligned, and the fluorescence intensities in the irradiated and unirradiated areas were measured using CellProfiler 4.0.7 image analysis software (Stirling et al., 2021). The relative intensity of the irradiated area was calculated by performing double normalization. After background subtraction, the intensity of the irradiated area was divided by that of the nucleus, and then the intensity ratio was divided by the average ratio before irradiation.

For FRAP (Arimura et al., 2013), 10 images were collected (2.0% 488-nm laser transmission; 128 × 24 pixels; pinhole 800 μm; 10× zoom), a 1.6 μm diameter circle area was bleached (100% 488-nm laser transmission; 31 ms), and another 90 images were collected without intervals.

To compare Ku80 dynamics with ATM and γ-H2AX, 11-4 cells that expressed EGFP-Ku80 and AT5BIVA cells that expressed EGFP-ATM were sensitized with Hoechst 33342 at 0.8 μM for 1 h before performing the laser microirradiation assay (512 × 512 pixels).

For the overnight observation to study the changes in PCNA distribution at different cell cycle phases, 11-4 and AT5BIVA cells expressing mCherry-PCNA were plated on a 35-mm glass-bottom dish. Cells were observed with a spinning disk confocal microscope (CSU-W1; Yokogawa and Ti-E; Nikon) with a PlanApo VC 100× (NA 1.40) oil-immersion objective lens equipped with an electron-multiplying charge-coupled device (iXon+; Andor) and a 488-nm laser (Nikon; LU-N4) at 37°C under 5% CO2. The images were captured with the NIS-Elements analysis software ver. 5.1 (Nikon).

### Inhibitor treatment and immunofluorescence

The inhibitors against ATR (AZ20), ATM (KU55933), and DNA-PK (NU7441) were purchased from Tocris Bioscience and dissolved in dimethyl sulfoxide (DMSO; Nacalai Tesque). To optimize the concentration of each inhibitor, cells plated in EZVIEW™ Glass Bottom Culture Plates LB (24 well; AGC Techno Glass) a day before were treated with each inhibitor at 10, 5, and 2.5 μM or with DMSO alone, simultaneously with etoposide (ETP; Sigma-Aldrich) at 20 μg/mL for 1 h at 37°C. The following procedures were performed at room temperature. Cells were fixed with 4% paraformaldehyde (PFA; Electron Microscopy Sciences) in 250-mM HEPES-NaOH (pH 7.4) containing 0.1% Triton X-100 (Nacalai Tesque) for 5 min at room temperature, washed with Dulbecco’s Phosphate Buffered Saline, calcium- and magnesium-free (PBS; Fujifilm Wako Chemicals), and permeabilized using 1% Triton X-100 for 20 min with gentle shaking. The cells were washed with PBS, incubated in blocking solution (Blocking-One P; Nacalai Tesque) for 20 min with gentle shaking, and washed with PBS. The cells were stained with γ-H2AX antibody (2 μg/mL) for 1 h with gentle shaking, washed with PBS three times, and stained with goat anti-mouse IgG (0.5 μg/mL; Jackson ImmunoResearch) conjugated with Alexa Fluor 488 (Thermo Fisher Scientific) and Hoechst 33342 (0.1 μg/mL; Nacalai Tesque) for 1 h with gentle shaking. The cells were washed with PBS before observation using a wide-field fluorescence microscope (Ti-E; Nikon) under the operation of NIS-Elements version 3.0 (Nikon) with a Plan Apo 40× (NA 0.95) objective lens, an electron-multiplying charge-coupled device (EM-CCD; iXon+; Andor; normal mode; gain ×5.1), an LF488-A filter set (Semrock), and a 75-W Xenon lamp as a light source.

MOF-knockdown cells were plated in a μ-Slide 8 well (ibidi) 1 day before ETP treatment (20 μg/mL) for 20 min at 37°C. The cells were fixed, permeabilized, and blocked as above before staining with fluorescence dye-labeled primary antibodies. MOF-knockdown cells that express mCherry-PCNA were stained Alexa Fluor 488-conjugated γ-H2AX and Cy5-conjugated H4K16ac (Hayashi–Takanaka et al., 2015) overnight at 4°C with gentle shaking. MOF-knockdown AT5BIVA cells that express EGFP-ATM were stained with Cy5-conjugated γ-H2AX and Cy3-conjugated H4K16ac overnight in Can-Get-Signal® Immunostain Immunoreaction Enhancer Solution B (Toyobo). The cells were washed with PBS, stained with Hoechst 33342 (0.1 μg/mL) for 1 h at room temperature with gentle shaking, and washed again with PBS before observation using a spinning-disk confocal microscope (CSU-W1; Yokogawa and Ti-E; Nikon) under the operation of NIS-Elements version 5.1 (Nikon) with a PlanApo 40× (NA 0.95) objective lens, an EM-CCD (iXon+; Andor; EM gain 300; gain ×5.1) and a 405/488/561/647 laser system (LU-N4; Nikon).

The image analysis was performed using the NIS-elements Analysis software ver. 5.1 (Nikon) by which nuclear areas were automatically defined by thresholding using Hoechst 33342 signals. The fluorescence intensity of each channel in the individual nucleus was then measured.

### Western blotting

Cell lysates were prepared by collecting the cultured cells by trypsinization, washing with cold PBS (Takara), and resuspension in lysis buffer [150 mM NaCl, 1% Triton X-100, 0.5% sodium deoxycholate (Fujifilm Wako Chemicals), 50 mM Tris-HCl pH8.0 (Nacalai Tesque)]. The protein concentration was measured by using the Protein Assay BCA Kit (Fujifilm Wako Chemicals) and bovine serum albumin as the standard according to the manufacturer’s instructions. Each sample was mixed with a sample-loading buffer (125 mM Tris-HCl, pH 6.8, 20% glycerol (Fujifilm Wako Chemicals), 4% sodium dodecyl sulfate (SDS; Fujifilm Wako Chemicals), 0.01% bromophenol blue (Fujifilm Wako Chemicals), and 10% dithiothreitol (Fujifilm Wako Chemicals)), heated at 95°C for 10 min. Then, 5–15 μL of each sample was separated on 7.5% (for ATM and RNA polymerase II) or 15% (for MOF, GAPDH, H4K16ac, and H4) polyacrylamide gels (SuperSep™ Ace, 17 well pre-cast; Fujifilm Wako Chemicals). The proteins on the gels were transferred to FluoroTrans W PVDF Transfer Membranes (Pall; 90 minutes; 170 mA constant for a 9 cm × 9 cm membrane) using EzFastBlot (Atto) as a transfer buffer. The membranes were blocked with Blocking One (Nacalai Tesque) for 30 minutes with gentle shaking. After washing with TBS-T (20 mM Tris-HCl, pH 8.0, 150 mM NaCl, 0.02% Tween 20), the membranes were incubated with the primary antibody, rabbit anti-ATM (1:1000; Abcam; Y170; a gift from S. Tashiro), rabbit anti-MOF/MYST1 (1:1000; Bethyl Laboratories), mouse anti-H4K16ac (1:1000; CMA416) (Hayashi–Takanaka et al., 2015), mouse anti-RNA pol II CTD (1:1000) (Stasevich et al., 2014), and mouse anti-GAPDH (1:1000; Santa Cruz Biotechnology; 6C5) diluted in Can-Get-Signal® Solution 1 (Toyobo) overnight at 4°C. After washing the membranes with TBS-T three times, the membranes were incubated with horseradish peroxide-conjugated goat anti-mouse IgG (H+L) (1:2000; Jackson ImmunoResearch) or goat anti-rabbit IgG (H+L) (1:2000; Jackson ImmunoResearch) diluted in Can-Get-Signal® Solution 2 (Toyobo) for 1 h at room temperature. The membranes were then washed with TBS-T. For chemiluminescence detection using a gel imaging system (LuminoGraph II, Atto), Western Lightning® Plus-ECL (PerkinElmer) and ImmunoStar® LD (Fujifilm Wako Chemicals) were used for RNA polymerase II and other proteins, respectively. For detecting total H4, WB Stripping Solution Strong (Nacalai Tesque) was used to strip the H4K16ac antibody before reprobing with a pan H4 antibody that binds to H4 regardless of the modification states (1:1000; CMA401) (Hayashi–Takanaka et al., 2015).

### Cell permeabilization

To extract proteins that freely diffuse in cells, the cells were permeabilized as described previously (Kimura et al., 2006). Cells that had been plated on a 35-mm glass-bottom dish 1 day before were chilled on ice, and washed twice with ice-cold PBF (100 mM potassium acetate, 30 mM KCl, 10 mM Na2HPO4, 1 mM dithiothreitol, 1 mM MgCl2, 1 mM adenosine triphosphate (Thermo Fisher Scientific), and 5% Ficoll (Nacalai Tesque)) followed by incubation with ice-cold PBF containing 0.1% TritonX-100 for 5 minutes. The cells were washed twice with cold PBF and incubated with Cy5-conjugated γ-H2AX Fab and Alexa Fluor 488-conjugated H4K20me2 Fab in PBF for 3–4 h on ice. Laser irradiation assay and observation were performed at 29°C using a confocal microscope (FV-1000), as described above.

### Statistical analysis

Student’s *t*-test (two-tailed) was performed using IBM SPSS Statistics for Windows, version 22 (IBM Corp.) Statistical significance was defined as *p* < 0.05.

## Acknowledgement

The authors are grateful to Satoshi Tashiro for AT5BIVA and 11-4 cells, and anti-ATM antibodies, Tsuyoshi Ikura for the ATM construct, Harumi Ueno and Yuko Sato for constructing the expression plasmid of EGFP-ATM, Takeshi Shimi for instructing the lentiviral shRNA infection experiment, members of Kimura lab for helpful discussion and suggestion, and the Biomaterials Analysis Division, Tokyo Institute of Technology for DNA sequencing. A part of this study was conducted through the Joint Usage/Research Center Program of the Radiation Biology Center, Kyoto University.

## Author Contributions

Conceptualization: W.T. and H.K.; Methodology: W.T. and H.K; Validation: W.T.; Formal analysis: W.T.; Investigation: W.T.; Writing - original draft: W.T.; Writing - review & editing: H.K.; Supervision: H.K.; Funding acquisition: H.K.

## Funding supports

This work was supported by the Ministry of Education, Culture, Sports, Science and Technology/Japan Society for the Promotion of Science KAKENHI (JP18H05527 and JP21H04764) and Japan Agency for Medical Research and Development (AMED) Basis for Supporting Innovative Drug Discovery and Life Science Research (BINDS) 21am0101105j0005 to H. Kimura.

## Supplemental figure legends

**Fig. S1.**
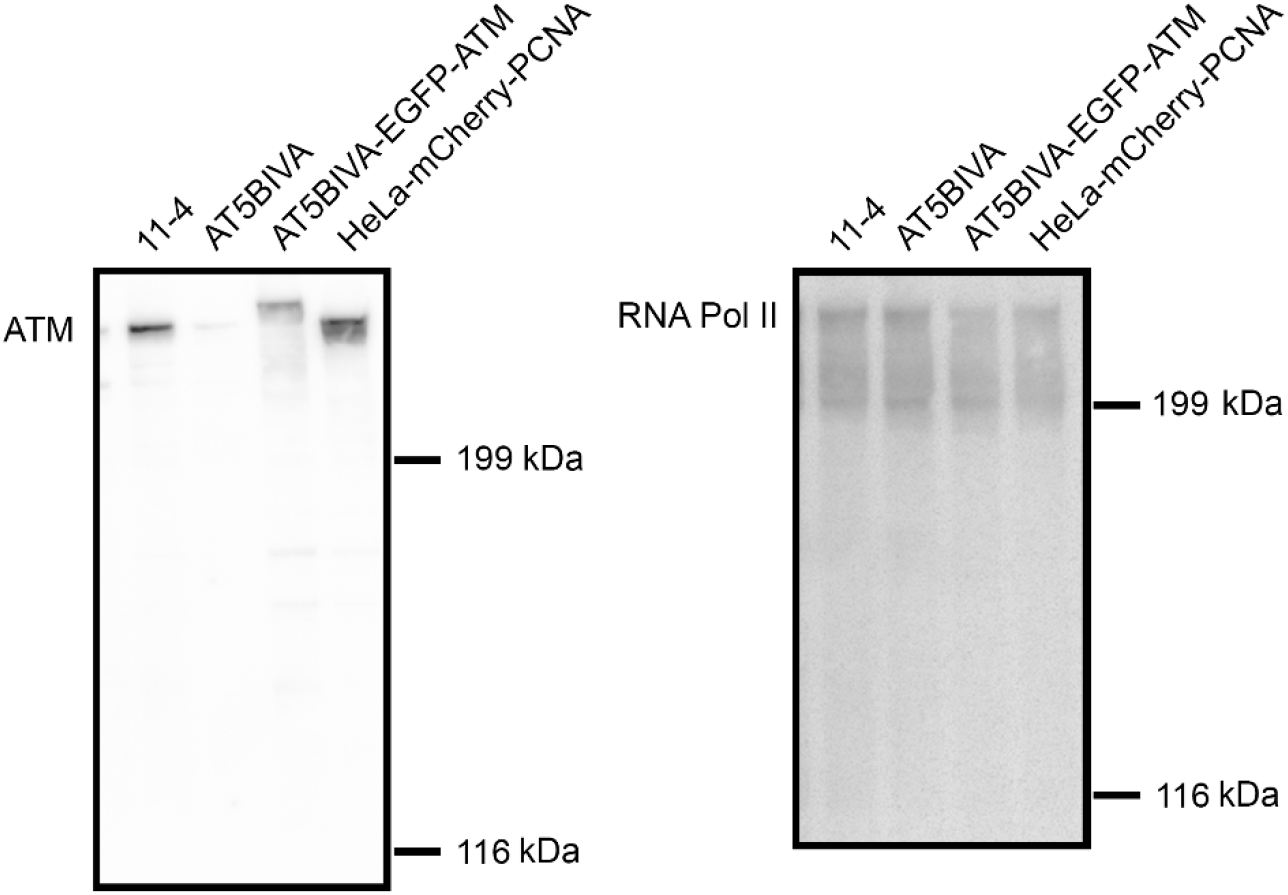
ATM expression in different cell lines. Cell lysates were separated on 7.5% SDS-polyacrylamide gels, transferred to membranes, and blotted with antibodies specific for ATM (left) and RNA polymerase II (right). Positions of size standards are indicated on the right.

**Fig. S2.**
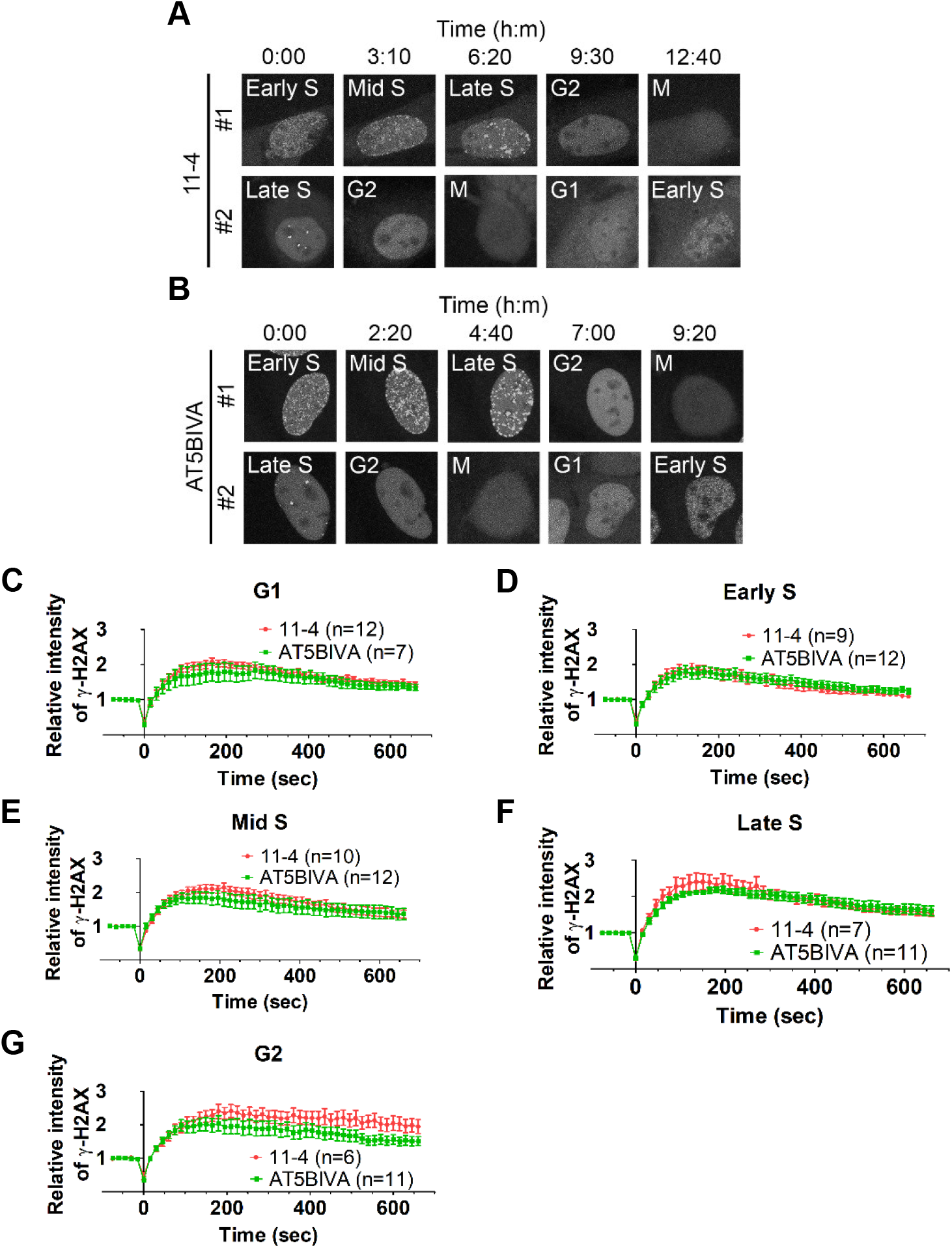
Accumulation kinetics of γ-H2AX in cells at different cell-cycle phases. To judge the cell-cycle phase of individual cells, mCherry-PCNA was expressed in 11-4 and AT5BIVA cells. (A and B) Time-lapse images of mCherry-PCNA in 11-4 (A) and AT5BIVA cells. In G1 cells, mCherry-PCNA distributed in both the nucleus and cytoplasm. Numerous foci of mCherry-PCNA are detected in the early-S phase and become bigger and fewer as the cell progresses through the mid-to late-S phase. mCherry-PCNA distributed throughout the nucleus except the nucleoli in the G2 phase. (C–G) Accumulation kinetics of γ-H2AX in G1 (C), early-S (D), mid-S (E), late-S (F), and G2 (G) phases. Graphs indicate the mean ± SEM with the number of cells (n). Scale bar: 5 μm.

**Fig. S3.**
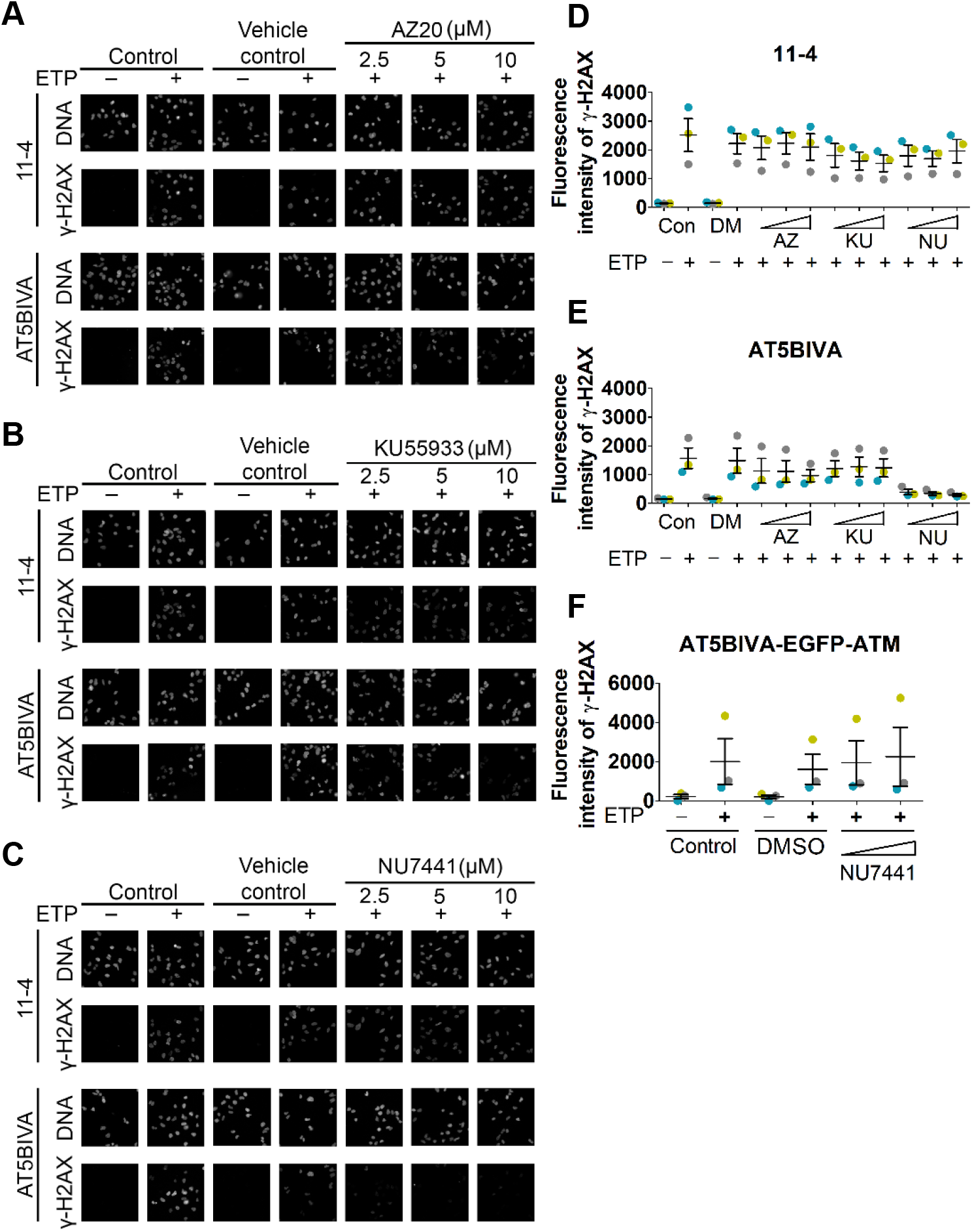
Optimizing the concentration of ATR, ATM, and DNA-PK inhibitor. (A–E) Treatment of 11-4 and AT5BIVA cells with AZ20, an ATR inhibitor (A), KU55933, an ATM inhibitor (B), and NU7441, a DNA-PK inhibitor (C), at 0, 2.5, 5, and 10 μM without or with ETP at 20 μg/mL for 1 h. The cells were fixed and stained with γ-H2AX-specific antibody followed by Alexa Fluor 488-conjugated anti-mouse antibody. DNA was stained with Hoechst 33342. (A–C) Fluorescence images. (D and E) The γ-H2AX fluorescence intensities in individual nuclei of 11-4 (D) and AT5BIVA (E) cells were measured in ≥60 nuclei per experiment and the mean intensities were obtained. Means ± SEM from three independent experiments are plotted with each data point. (F) AT5BIVA cells expressing EGFP-ATM were treated with NU7441 at 2.5 and 5 μM simultaneously with ETP at 20 μg/mL for 1 h. The cells were fixed and stained with γ-H2AX-specific antibody followed by Cy3-conjugated anti-mouse antibody. Means ± SEM are plotted with each data point as in (D) and (E). Scale bar: 5 μm.

**Fig. S4.**
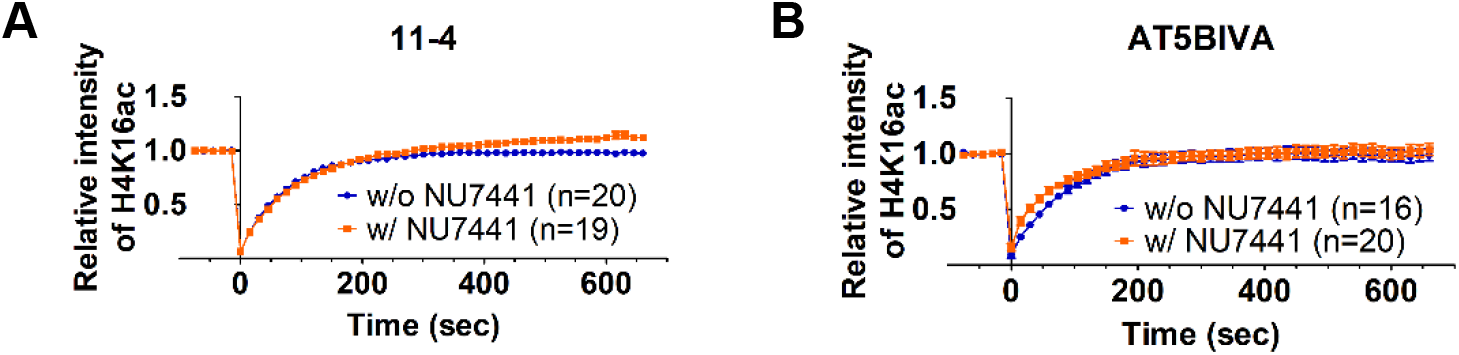
H4K16ac in 11-4 and AT5BIVA cells. Loading of 11-4 (A) and AT5BIVA (B) cells with H4K16ac Fab followed by treatment without or with 2.5 μM of NU7441 for ≥1 h before inducing DNA damage by laser irradiation. The Fab intensities in the irradiated areas were measured and plotted (mean ± SEM, with the number of cells, n).

**Fig. S5.**
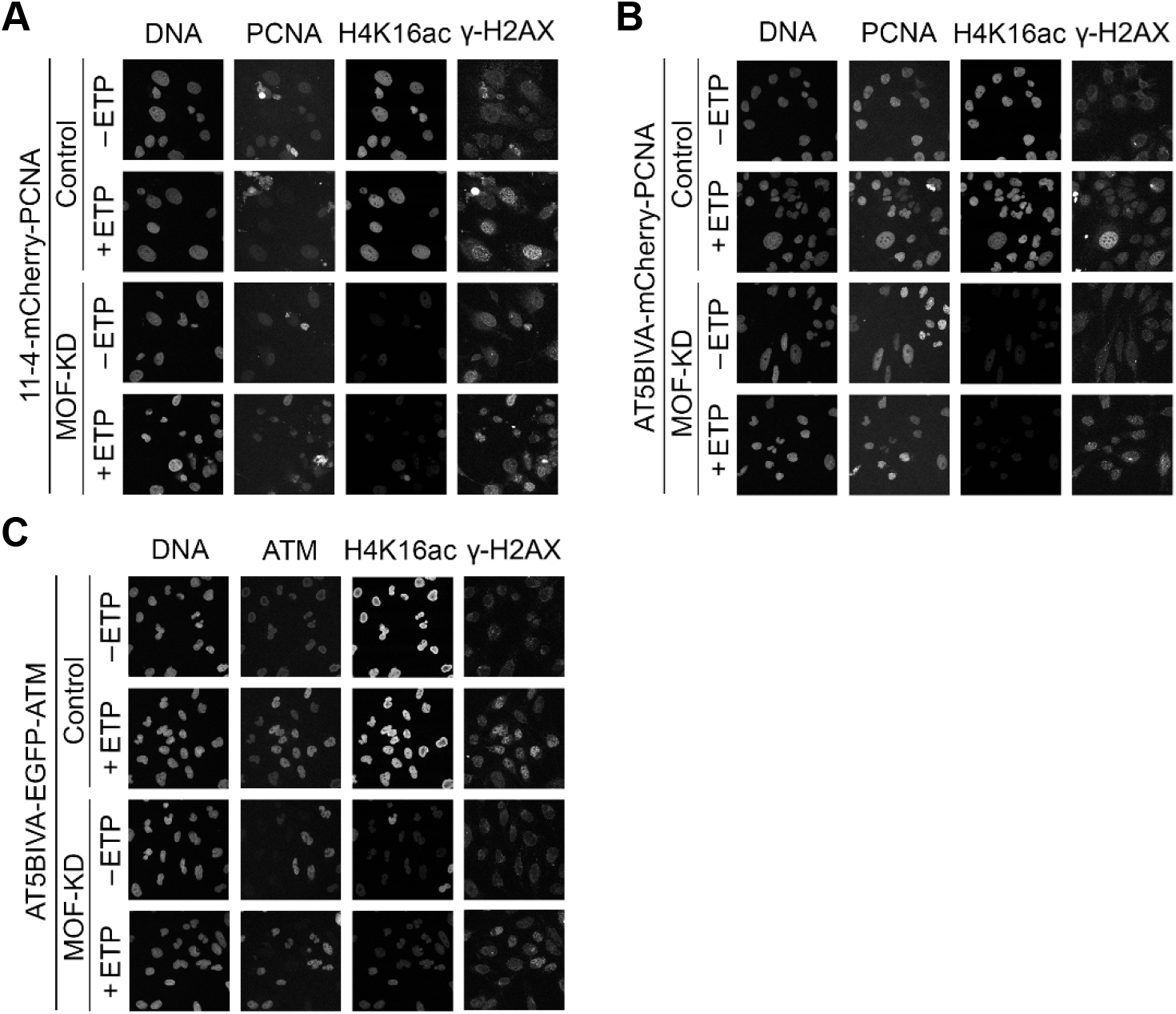
H4K16ac and γ-H2AX in MOF-knockdown cells. MOF was knocked down in 11-4 cells expressing mCherry-PCNA (A), AT5BIVA cells expressing mCherry-PCNA (B), and AT5BIVA cells expressing EGFP-ATM. The cells were treated with ETP at 20 μg/mL for 20 min to induce DNA damage, and fixed. The 11-4 and AT5BIVA cells expressing mCherry-PCNA were stained with Alexa Fluor 488-labeled γ-H2AX and Cy5-labeled H4K16ac antibodies. The AT5BIVA cells expressing EGFP-ATM were stained with Cy3-labeled H4K16ac and Cy5-labeled γ-H2AX antibodies. Scale bar: 5 μm.

**Fig. S6.**
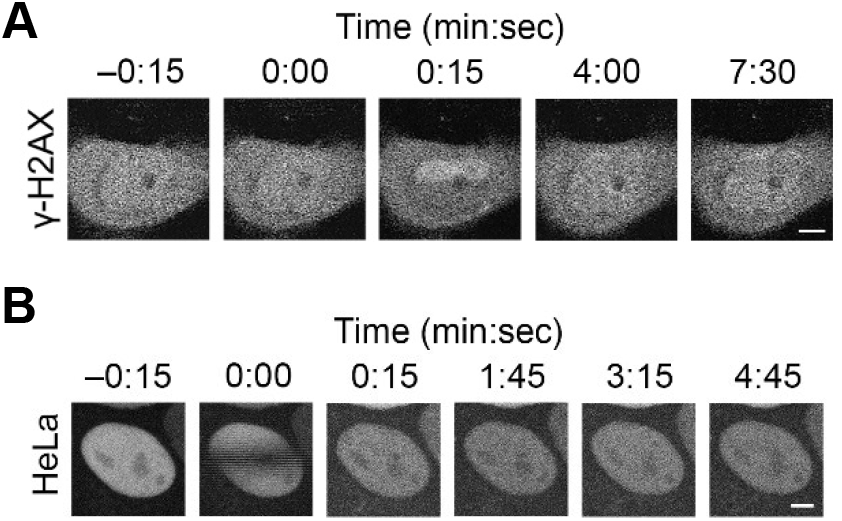
EGFP-Ku80 and EGFP-ATM in Hoechst 33342-sensitized cells. (A) Accumulation of γ-H2AX in AT5BIVA cells expressing EGFP-ATM that were sensitized with 0.8 μM Hoechst 33342 for 1 h before laser irradiation. (B) Accumulation of EGFP-Ku80 in HeLa cells without Hoechst 33342 sensitization. Scale bar: 5 μm.

